# New insights on folliculogenesis and follicular placentation in marine viviparous fish black rockfish (*Sebastes schlegelii*)

**DOI:** 10.1101/2021.01.20.427513

**Authors:** Xiaojie Xu, Qinghua Liu, Xueying Wang, Xin Qi, Li Zhou, Haoming Liu, Jun Li

**Affiliations:** The Key Laboratory of Experimental Marine Biology, Center for Ocean Mega-Science, Institute of Oceanology, Chinese Academy of Sciences, Qingdao, China; Laboratory for Marine Biology and Biotechnology, Qingdao National Laboratory for Marine Science and Technology, Qingdao, China; University of Chinese Academy of Sciences, Beijing, China; Key Laboratory of Mariculture (Ocean University of China), Ministry of Education, Ocean University of China, Qingdao, China; Weihai Shenghang Aquatic Science and Technology Co., LTD. Weihai, China; Fisheries Research Institute of Huancui District, Weihai China

**Keywords:** Folliculogenesis, Follicular placentation, Viviparous fish, *Sebastes schlegelii*

## Abstract

In viviparous fish, a considerable degree of variation in placental structures have been described. However, no distinct structures are reported in Scorpaenidae. In this study, we demonstrate a new type of folliculogenesis and follicular placentation in *Sebastes schlegelii*. Before copulation, the germinal epithelium gradually surrounds the oocytes and develops to individually follicles with a stalk-like structure hanging on the ovigerous lamella, which ensures each follicle have access to spermatozoa after copulation. From stage V to early gestation, the *cyp17-I* highly expressed accompanied by *cyp19a1a* signals disappearance, and 11-ketotestosterone level keeps rising and peaks at blastula stage, while 17β-estradiol declines to the bottom. Meanwhile, the theca cells rapidly proliferate and invade outwards forming a highly hypertrophied and folded microvillous placenta. This unbalance of hormone might be an important factor driving the theca cells proliferation and invasion. Additionally, some conserved genes related to mammalian placentation are significantly high expression in follicular placenta suggesting the high convergence in vertebrate placenta evolution.

## 1. Introduction

Viviparity reproduction is a wide-spread reproductive strategy (*Kalinka, 2015*). It earliest arose among fishes, and it occurred in most vertebrates, including most cartilaginous fishes, several clades of bony fishes, amphibians, reptiles and most mammals (*Wourms and Callard, 1992; Wourms, 1993; Blackburn and Daniel, 2015*). The developing embryos were retained within the parental body, supported by its own reserved yolk or provision of maternally derived nutrients, and leaded to release of live offspring instead of egg. Over 500 species of teleost fish in 14 families have been identified as viviparity (*Kunz et al., 2004; Blackburn, 2005*).

For viviparity, initial steps in the evolution of live-bearing from egg-laying must involve a shift from external to internal fertilization. The male transferred the sperm to the female gonaduct and fertilized the eggs(*Carcupino et al., 2002; Kwan et al., 2015*). Therefore, morphological and physiological adaptations of the ovary to facilitate the maternal-embryo interaction is an obligatory aspect of viviparity (*Roberts et al., 2016*). In fish, since the first “follicular pseudoplacenta” or a placental analogs in poeciliidae was described (*Turner, 1940; Knight et al., 1985*), a considerable degree of variation in placental structures were reported, including umbilical cord in shark (*Buddle et al., 2018*), brood pouch of the male in sygnathid (*Laksanawimol et al., 2006; Stolting and Wilson 2007*) or the ovarian gestation in Zoarcidae (*Larsson et al., 2002*), Cyprinodontiformes and Sebastinae (*Blackburn 2005; Reznick et al 2002, Knight et al., 1985, Kwan et al., 2015*). Intraovarian gestation is unique among vertebrates (*Wourms et al., 1988; Schindler and Hamlett, 1993*), for lacking Mullerian ducts from which oviducts develop in other vertebrates (*Turner 1947; Wake 1985; Campuzano Caballero and Uribe, 2014*).

Placenta is a transient organ to facilitate the nutrition, gases and waste exchange and to regulate maternal-fetal interactions often through hormone production (*Van Dyke et al., 2014; Burton and Jauniaux, 2015*). The evolution of a novel organ typically involves both functional innovations and a novel structure which is associated with this function. In both *Poeciliopsis retropinna* and *P. turneri* (*Wourms, 1981; Wourms et al., 1988; Kwan et al., 2015; Guernsey et al., 2020*), the inner surface of the maternal follicular epithelium was highly hypertrophied and extensively folded (*Grove and Wourms, 1994; Kwan et al., 2015*). In Goodeidnae, at mid to late gestation stages, the embryos moved from follicle to the ovarian lumen and developed a trophotaeniae (*Knight et al., 1985; Lombardi and Wourms, 1985; Wourms and Callard, 1992; Uribe et al., 2018*).

In Scorpaenidae, four genera (Sebastes, Sebasticus, Helicolenus and Hozukius) also belong to the marine viviparity (*Turner, 1947*), but no obviously developed structures were observed (*Moser, 1967*). Black rockfish (*Sebastes schlegelii*) is an important comercial marine species which inhabits in North China, Korea, and Japan. The female and male copulate in November, while fertilization occurs in the next March. After almost 2 months of pregnancy in the ovary, the offspring are released into the sea (*Shi et al., 2011; Mori et al., 2003*). Usually, the enormous fecundity was diminished to adapt internal fertilization and gestation (*Grier et al., 2005*). Interestingly, the fecundity of black rockfish is high and comparable to that of oviparous fishes ranging from 35,000 to 472,000 (*Nakagawa 1998, 1999, 2000, 2001; Nakagawa et al. 2002*). Therefore, it raises an intriguing question, how does the black rockfish modify the morphology, physiology and gene expression profile to adapt this reproductive strategy. In this study, we investigated the developmental process of oogenesis and gestation and found that the folliculogeneis of black rockfish is different from oviparous species and other documented viviparous placentas. When the oocytes developed to early secondary growth (SGe) stage, the follicles were surrounded by the germinal epithelium with stalk-like structures attached the ovigerous lamella. The results from *in situ* hybridization (ISH), steroid hormone changes and transcriptome indicated the dramatical expression of cytochrome P450c17 (*cyp17-I*) from full secondary growth (SGf) to blastula stage gave rise to the theca cells rapidly proliferation, migration and invasion into the stroma and formed a new type of follicular placenta. Additionally, the closely associated genes with mammalian placentation including *HLA-E, laminin α4* (*lama4*), *placenta special gene 8* (*plac8*), *trophoblast glycoprotein* (*tpbg*), placenta growth factor (*plgf*) expressed strongly throughout placentation suggesting these high conserved genes were convergent in the vertebrate placentation.

## 2. Results

### 2.1 Oogenesis and gestation of *Sebastes schlegelii*

Oogenesis and embryonic development of black rockfish are shown in figure1. For both black rockfish and turbot, the germinal epithelium bordered the various developing stage oocytes and formed the ovigerous lamella. But for black rockfish, it contained much more richer stroma in comparision with turbot (Fig. 1a-c). When the oocytes developed into the SGe stage, they were surrounded by the germinal epithelium, but still remained attaching to the stroma through a stalk-like structure. At this stage, the female and male copulated, and the male transferred spermatozoa to the ovarian cavity (Fig. 1c-c2). Numerous spermatozoa scattered in the ovarian lumen immediately after copulation, and stored in the crypt between the epithelium cells and theca layer or at the folds outside the follicles (Fig. 1d-e). At SGf stage, the vitellogenesis finished, oocytes entered into maturation and follicle layers broke down (Fig. 1f). Then the eggs fertilized with the spermatozoa which hidden in the crypt or folds before. At the same time, the granulosa detached from the zona pellucida (ZP), and mixed with the surrounding theca layers forming a barrier. After that, they rapidly migrated and invaded into the surrounding tissues and formed a follicular placenta to support the embryos development (Fig. 1g).

**Figure 1.**
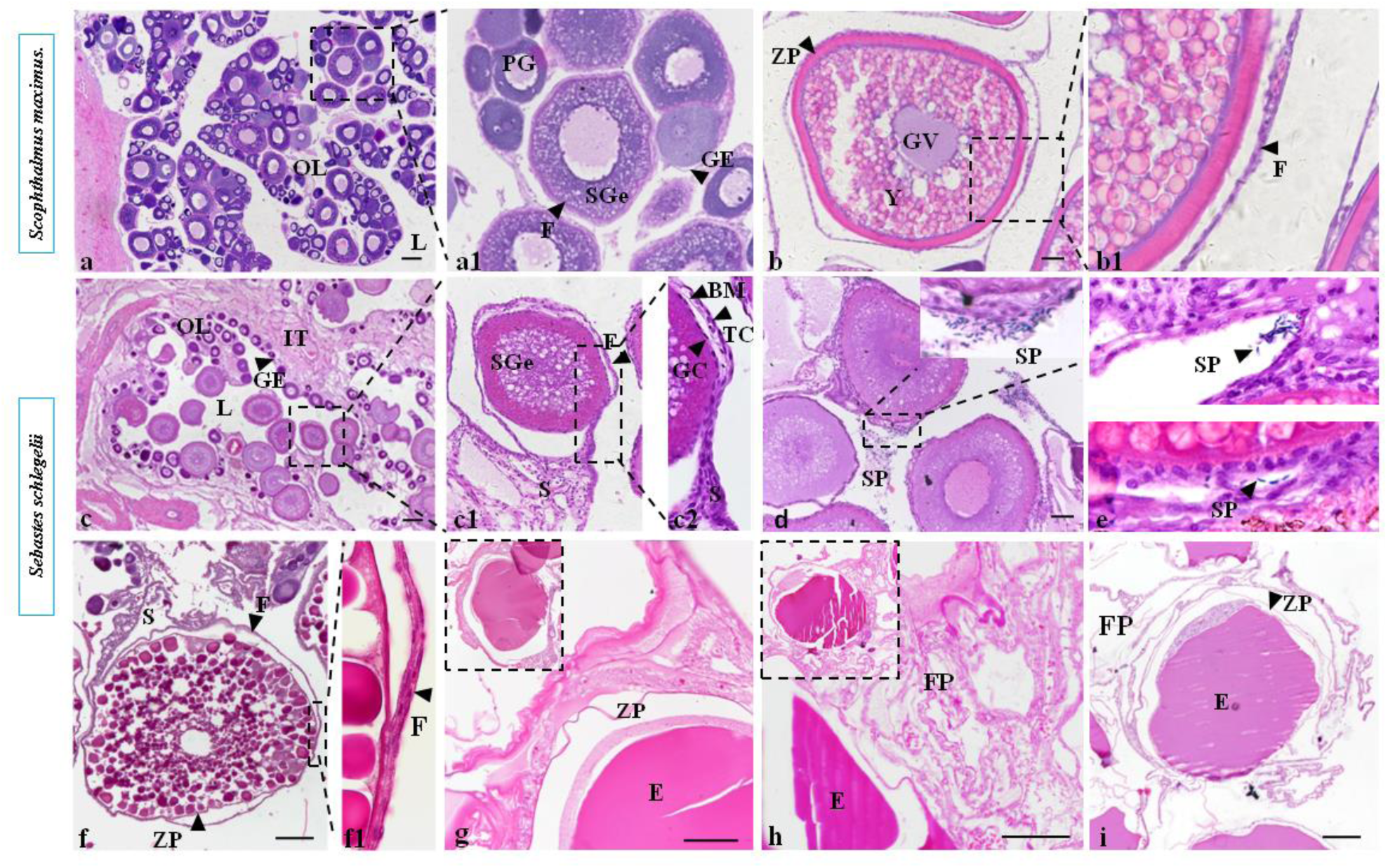
Oogenesis and embryonic development of black rockfish compared with turbot. Numerous primary growth oocytes and early secondary growth (SGe) oocytes surrounded by follicle cells in the stage III ovary of turbot (a, a1). Full secondary growth (SGf) oocytes surrounded by a thick zona pellucida (ZP) and thin follicle cells (F) in the stage V ovary of turbot (b, b1). Numerous primary growth oocytes and early secondary growth (SGe) oocytes surrounded by follicle cells in the stage III ovary of black rockfish (c, c1). Numerous spermatozoa of scatter in the ovarian lumen outside of the follicles in the stage III ovary of black rockfish (d, d1). Numerous spermatozoa hide in the crypt of the stromal cells or the folds outside of the follicles in the stage IV ovary of black rockfish (e). SGf oocytes are surround by a thin ZP and follicular layers in the stage V ovary of black rockfish (f). At cleavage stage, the granulosa cells have detached from the oocyte and the follicular layers (granulosa layer,theca layer and basement membrane) mixed with the surrounding epithelium and stroma cells and formed follicular placenta (g). Follicular placenta structure became highly hypertrophied, extensively folded at blastula stage (h). Follicular placenta became more loose at gastrulae stage (i). L, lumen; OL, ovigerous lamella; IT, intersticial tissue; PG, follicles with primary growth; GE, germinal epithelium; TC, theca cells; GC, granulosa cells; BM, basement membrane; YG, yolk globule; BV, blood vessel; ZP, zona pellucida; S, stroma; SGf, full secondary growth; SGe, early secondary growth; F, follicle layers; FP, follicular placenta; E, embryo. Scale bars, 200μm

### 2.2 Characterization of *HLA-E*

The open-reading frame of black rockfish *HLA-E* is 1170bp. The deduced *HLA-E* protein is composed of 389 amino acids. The results of conserved domain showed that black rockfish had the same conserved domains as humans (Fig. 2a). Since *HLA-E* belongs to the major histocompatibility complex class I family (MHC-I), the phylogenetic analysis was conducted using the predicted amino acid sequences to analyze the evolutionary relationship of the major histocompatibility complex classIfamily (MHC-I). The MHC-I are divided into two main groups. *Sebastes schlegelii, Oplegnathus fasciatus, Perca flavescens, Epinephelus lanceolatus, Homo sapiens* and *Pongo abelii* clustered in one of the subbranches. These results showed that *HLA-E* gene of *Sebastes schlegelii* was homologous to *Homo sapiens* (Fig. 2b)

**Figure 2.**
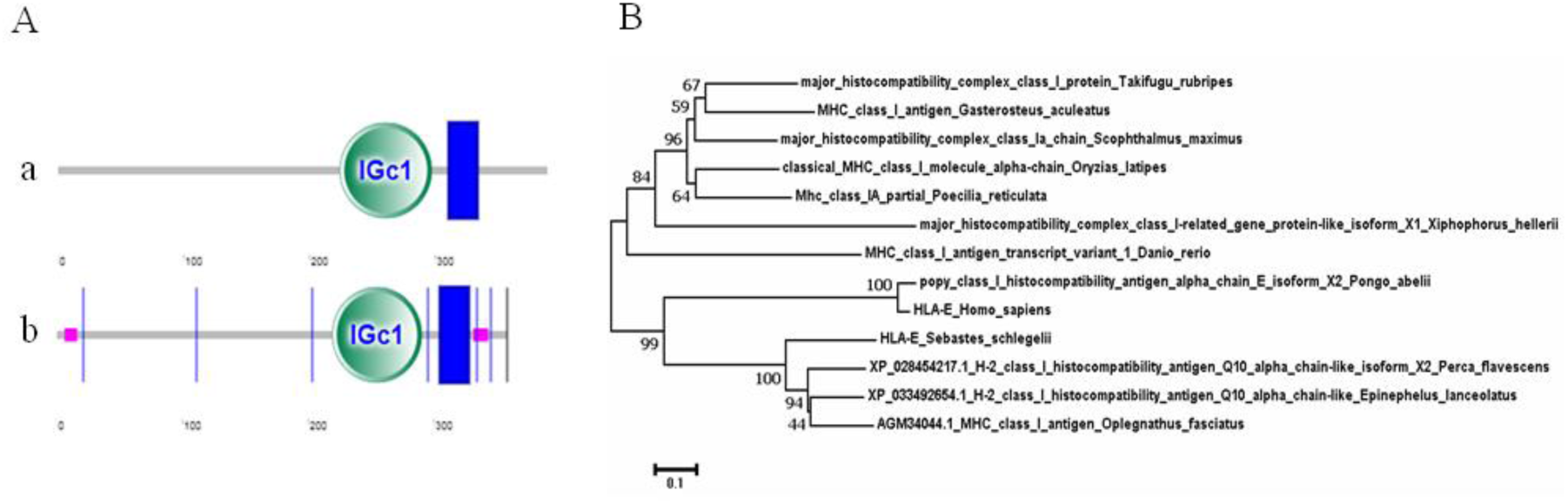
The conserved domains and phylogenetic tree of HLA-E in black rockfish A. The HLA-E of black rockfish has the same conserved domains as humans. Conserved domains of black rockfish (a). Conserved domains of human (b). B. The phylogenetic tree of the major histocompatibility complexclass I family (MHC-I) includes black rockfish and other vertebrates using predicted amino acid sequences. The GenBank accession numbers are as follows: *Xiphophorus macufulls* H-2 class I histocompatibility antigen, alpha chain-like (XP_023201134.1), *Poecilia reticufull* PREDICTED: H-2 class I histocompatibility antigen, Q10 alpha chain-like isoform X1 (XP_008420844.1), *Homo sapiens* HLA-E (ARB08449.1), *Pongo abelii*popy class I histocompatibility antigen, alpha chain E isoform X2 (XP_024104292.1), *Danio rerio*MHC class I antigen transcript variant 1 (ALL98461.1), *Gasterosteus aculeatus* MHC class I antigen (ABN14357.1), *Scophthalmus maximus*major histocompatibility complex class Ia chain (ABM92962.1), *Takifugu rubripes* major histocompatibility complex class I protein (AAC41236.1), *Oryziaslatipes* classical MHC class I molecule, alpha-chain (BAJ07297.2), *Oplegnathus fasciatus* MHC class I antigen (AGM34044.1), *Perca flavescens*H-2 class I histocompatibility antigen, Q10 alpha chain-like isoform X2 (XP_028454217.1), *Epinepheluslanceolatus*H-2 class I histocompatibility antigen, Q10 alpha chain-like (XP_033492654.1).

### 2.3 The changes of hormone level and related genes expression during the process of ovarian development

Two-cell type model illustrating the interaction of granulosa layers and theca cells of the ovarian follicle in the biosynthesis of active steroid hormones in gonad are shown in figure 3A. In vertebrates, granulosa and theca cells in the follicles are responsible for the steroid hormone biosynthesis, including gonadal steroid hormones, progesterone, estradiol (*Lubzens et al., 2010, Sreenivasulu and Senthilkumaran, 2009*). After cholesterol is catalyzed by the cholesterol CYP11A1, pregnenolone and progesterone undergo 17a-hydroxylation and proceed down the C21, 17-hydroxy pathway to 17a-hydroxypregnenolone and 17a-hydroxyprogesterone, respectively. CYP17 has17a-hydroxylase and 17,20-lyase activity (*Miller et al., 1997; Kagawa, 2013*), and only specifically expresses in the specific steroid-production theca cells in the theca layers, which is critical not only for maintaining the structural integrity of the follicle but also for delivering nutrients to the avascular granulosa cell layer. And the theca-derived androgens are then converted to estradiol by the CYP19A1 enzyme in granulosa cells.

**Figure 3.**
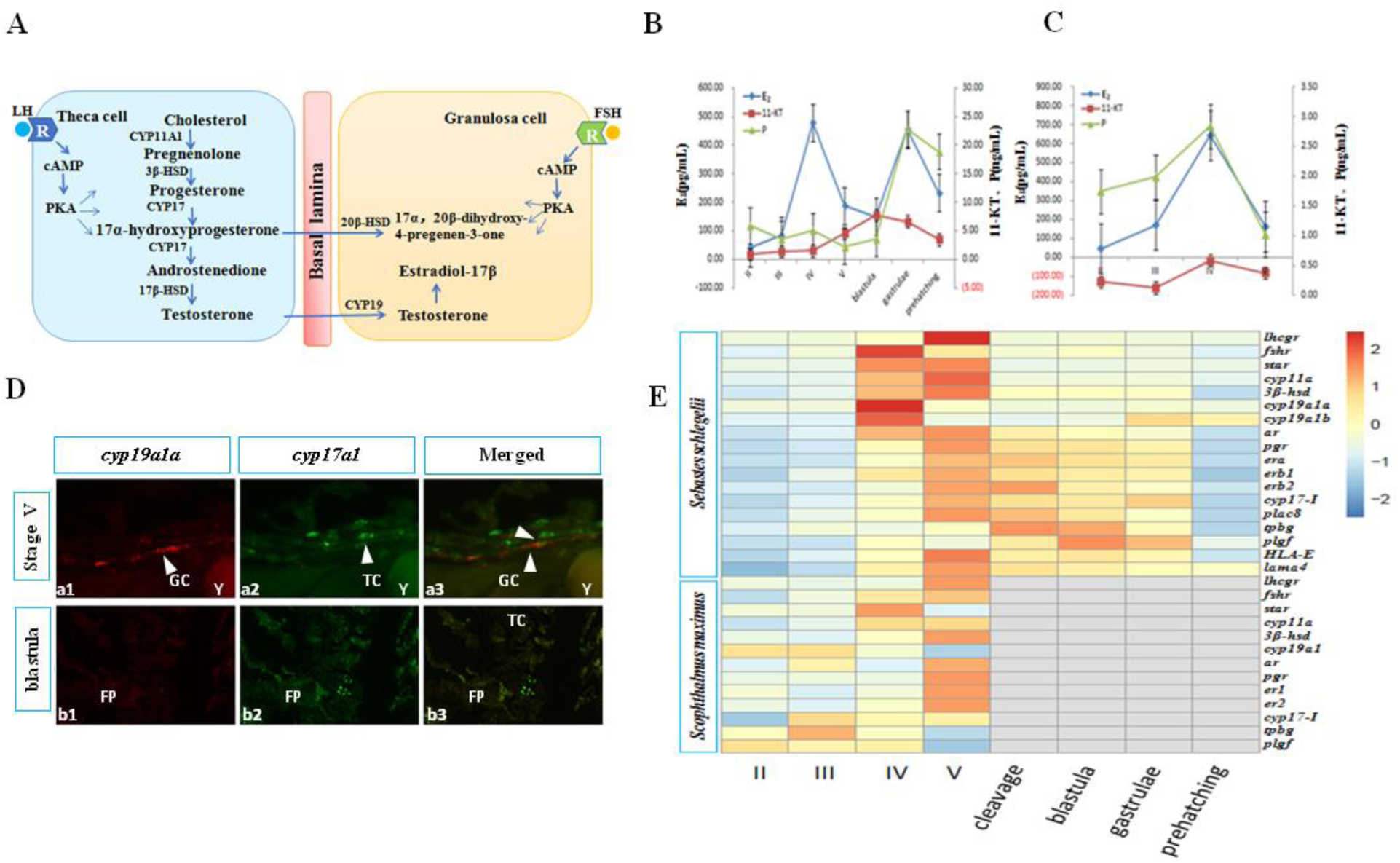
The changes of hormone level and related genes expression during the process of the ovarian development. A. Two-cell type model illustrates the Interaction of granulosa layers and theca cells of the ovarian follicle in the biosynthesis of active steroid hormones in gonad. Enzymes: P450scc (CYP11A1), P450 side-chain cleavage; P450c17(CYP17), 17-hydroxylase/C17-C20-lyase; 3b-HSD, 3b-hydroxysteroid dehydrogenase;17b-HSD, 17b-hydroxysteroid dehydrogenase; 20b-HSD, 20b-hydroxysteroid dehydrogenase; P450arom (CYP19A), P450 aromatase. B. Three steroid hormone changes during the oogenesis and placentation of black rockfish. 11-KT had been at a low level until the ovary developed to stage IV. When the ovary developed to stage IV, 11-KT level gradually rose and peaked at blastula stage. After that, it decreased slightly but still remained at a relatively high level throughout the pregnancy. The level of E2 increased significantly from stage III to stage V, and decreased at blastula stage. During pregnancy, E2 also maintained a high level, and peaked again during the gastrulae period. The level of P was low until blastula stage, it rapidly rose and remained a high level during the pregnancy period. C. Three steroid hormone changes during the oogenesis of turbot. The 11-KT,E2 and P level presented an upward trend,peaked stage IV and decreased from stage IV to stage V. D. The results of two-color fluorescence in situ hybridization of *cyp17-I* and *cyp19a1a* at SGf and blastula. The expression of *cyp17-I* with green and *cyp19a1a* with red at SGf of black rockfish (a1-a3). the expression of *cyp17-I* with green and *cyp19a1a* with red in blastula stage of black rockfish (b1-b3). E. Expression profile of some important genes during ovarian development at eight different development stages. The log ratio expression is indicated in a heat map. 11-KT, 11-ketotestosterone; E2, 17β-estradiol; P, progesterone; TC, theca cells; GC, granulosa cells; Y, yolk granules; FP, follicular placenta.

The results of steroid hormone of black rockfish and turbot are shown in figure 3B, figure 3C, respectively. The 11-ketotestosterone (11-KT) had been at a low level until stage IV, then gradually rose and peaked at blastula stage, and remained at a relatively high level throughout the pregnancy. In the process of vitellogenesis, the level of 17β-estradiol (E2) increased significantly and peaked at stage IV, then dramatically decreased from stage V to blastula stage. During mid to late pregnancy, E2 also maintained a high level, and peaked again during the gastrulae period. In the process of oogenesis, the level of progesterone (P) was low, while the level of progesterone rose rapidly and remained at a very high level at gestation stage (Fig. 3B). For turbot, the level of E2 presented an upward trend from stage II to stage IV, and decreased from stage IV to stage V. The change trend of progesterone and 11-KT were similar to that of E2 (Fig. 3C).

The results of two-color fluorescence *in situ* hybridization of *cyp17-I* and *cyp19a1a* at SGf and blastula stage are shown in figure 3D. When the oocytes developed to SGf stage, both *cyp17-I* with green signals and *cyp19a1a* with red signals expressed on theca cells and granulosa cells, respectively (Fig. 3a1-a3). However, when the embryo developed to the blastula stage, only *cyp17-I* showed signals on follicular placenta, and *cyp19a1a* had no obvious signals (Fig. 3b1-b3).

Expression profile of some important genes related to oogenesis and gestation at eight different stages are shown in figure 3E. For black rockfish luteinizing hormone/choriogonadotropin receptor (*lhcgr*), follicle stimulation hormone receptor (*fshr*), steroid acute regulatory protein (*star*), cholesterol side-chain cleavage enzyme (*cyp11a*), 3β-hydroxyl steroid dehydrogenases (*3β-hsd*), androgen receptor (*ar*) and cytochrome P450 aromatase (*cyp19a1a*) were highly expressed during oogenesis and weakly expressed during pregnancy. During oogenesis, these genes had the same expression trend in both black rockfish and turbot. The *cyp17-I*, progesterone receptor (*pgr*), estrogen receptor alpher (*era*), estrogen receptor beta1 (*erb1*), estrogen receptor beta2 (*erb2*), *plac8, tpbg, plgf, HLA-E* and *lama4* strongly expressed during pregnancy, especially *plac8, tpbg* and *plgf* in the early and mid pregnancy in black rockfish. *HLA-E, lama4, cyp19a1b, plac8* and *era* were not detected in turbot.

### 2.4 *Cyp17-I, cyp19a1a, fshr, lama4* and *HLA-E* mRNA location in gonad during the oogenesis

The expression of *cyp17-I, cyp19a1a, fshr, lama4* and *HLA-E* in gonad during oogenesis are shown in figure 4. When the oocytes developed to SGe stage, *cyp17-I* expressed on theca cells, and the signals gradually increased with the ovary development. When the oocytes developed to SGf stage, the signals of *cyp17-I* could be detected not only on theca cells, but throughout all the stromal cells around the oocytes, especially in stalk-like tissues (Fig. 4a1-a3). For *cyp19a1a*, it only expressed on granulosa cells and the signals got much stronger from SGe to late secondary growth (SGl) stage, and became very weak at SGf stage (Fig. 4b1-b3). While *fshr* signals presented similar position and change trend with *cyp19a1a* (Fig. 4c1-c3). *Lama4* signals could not be detected on theca cells until the oocytes developed to SGl stage, and significantly increased at SGf stage on theca cells as well as the stromal cells around the oocytes (Fig. 4d1-d3). Similarly, the signals of *HLA-E* expressed on theca cells and stromal cells around the oocytes, and gradually increased as the ovary developed (Fig. 4e1-e3).

**Figure 4.**
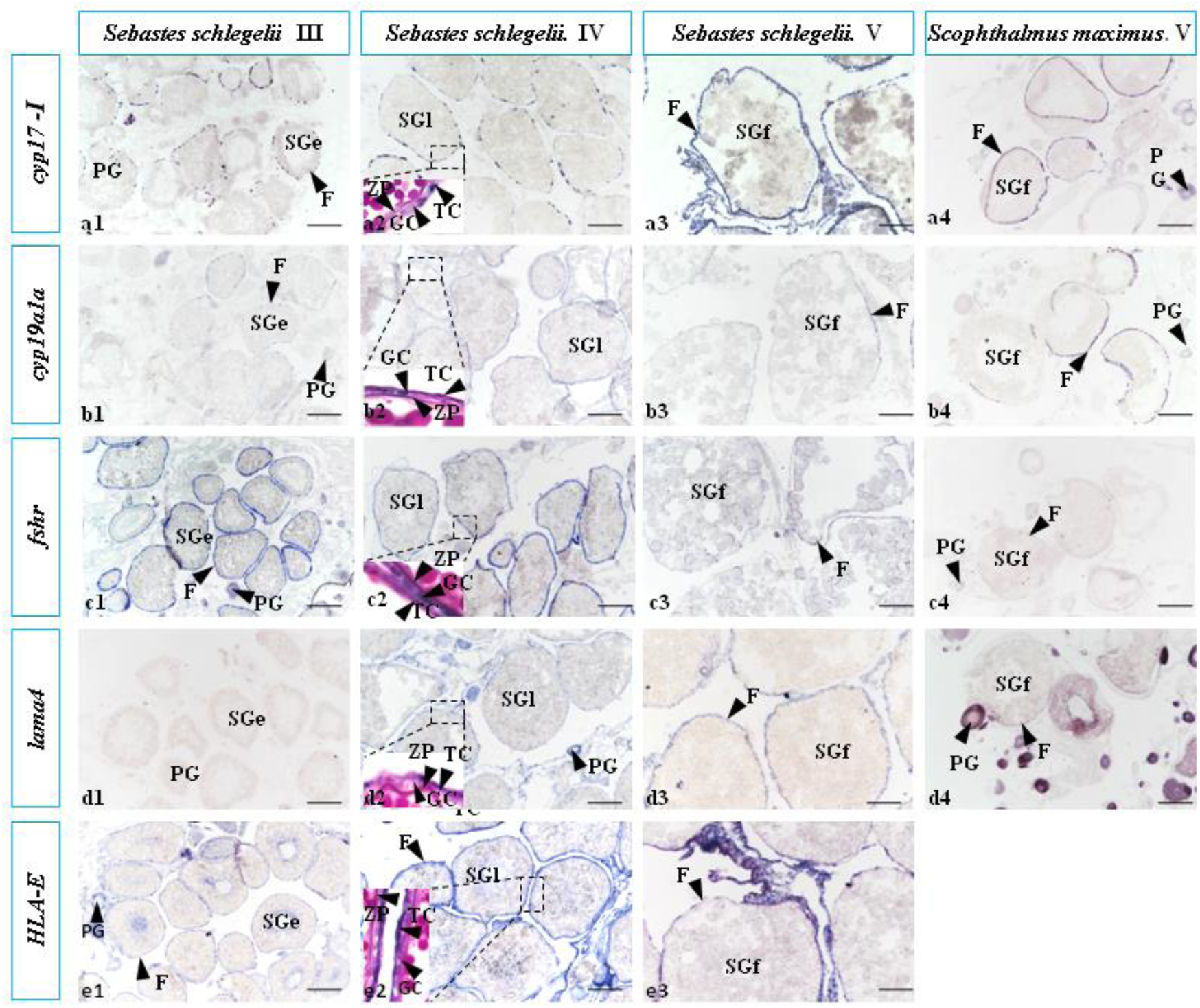
The expression of *cyp17-I, cyp19a1a, fshr, lama4* and *HLA-E* in gonad during the oogenesis. At SGe stage, *cyp17-I* expressed on theca cells, and the signals gradually increased with the ovary development. At SGf stage, the signals of *cyp17-I* could be detected not only on theca cells, but throughout all the stromal cells around the oocytes, especially in stalk-like tissues (a1-a3). For turbot,*cyp17-I* only expressed on theca cells, not on the stromal cells around the oocytes at SGf stage (a4). *Cyp19a1a* only expressed on granulosa cells, and the signals got stronger significantly from SGe to SGl stages, and became weak at SGf stage (b1-b3). For turbot, the signals of *cyp19a1a* expressed on granulosa cells (b4). *Fshr* only expressed on granulosa cells, and the signals got stronger significantly from SGe to SGl stage, but the signals became very weak at SGf stage in black rockfish (c1-c3). For turbot, the signals of *fshr* expressed on granulosa cells (c4). *Lama4* signals could not be detected on theca cells until the oocytes developed to SGl stage, and significantly increased at SGf stage on theca cells as well as the stromal cells around the oocytes in black rockfish (d1-d3). For turbot, *Lama4* had no signal at SGf stage (d4). *HLA-E* signals could be detected on theca cells and stromal cells from SGe stage, and significantly increased at SGf stage (e1-e3). TC, theca cells; GC, granulosa cells; PG, follicles with primary growth; ZP, zona pellucida; SGe, early secondary growth; SGl, late secondary growth; SGf, full secondary growth; F, follicle layers. Scale bars, 200μm.

For turbot, *cyp17-I* was also detected on theca cells, but the difference was that *cyp17-I* of turbot only expressed on theca cells, not on the stromal cells around the oocytes at SGf stage (Fig. 4a4). The expression of *cyp19a1a* and *fshr* of turbot was similar to black rockfish that the signals expressed on granulosa cells (Fig. 4b4, c4). *Lama4* had no signal at SGf stage, which was completely different from black rockfish during this period (Fig. 4d4).

### 2.5 *Cyp17-I, cyp19, fshr, lama4* and *HLA-E* mRNA expression in gonad during the pregnancy

The expression of *cyp17-I, cyp19a1a, fshr, lama4* and *HLA-E* during pregnancy are shown in figure 5. *Cyp17-I* signals presented strong on the follicular placenta at blastula and gastrulae stage, and disappeared until prehatching stage (Fig. 5a1-a3). *HLA-E* had the similar expression pattern with *cyp17-I* during gestation (Fig. 5e1-e3), while *fshr* and *cyp19a1a* had no obvious signals during this period (Fig. 5b1-c3). *Lama4* expressed on the follicular placenta during the whole pregnancy period, the signals were strong at blastula and gastrulae stage, and became weak at the prehatching stage (Fig. 5d1-d3). For the isolated females, no obvious follicular placenta structure was observed in the ovary, and only stromal cells, vascular structure and non-cellular structure surrounded the unfertilized eggs (Fig. 6a). And *lama4, cyp17-I, cyp19a1a* and *fshr* did not show any signals, except for *HLA-E*(Fig. 6b-f).

**Figure 5.**
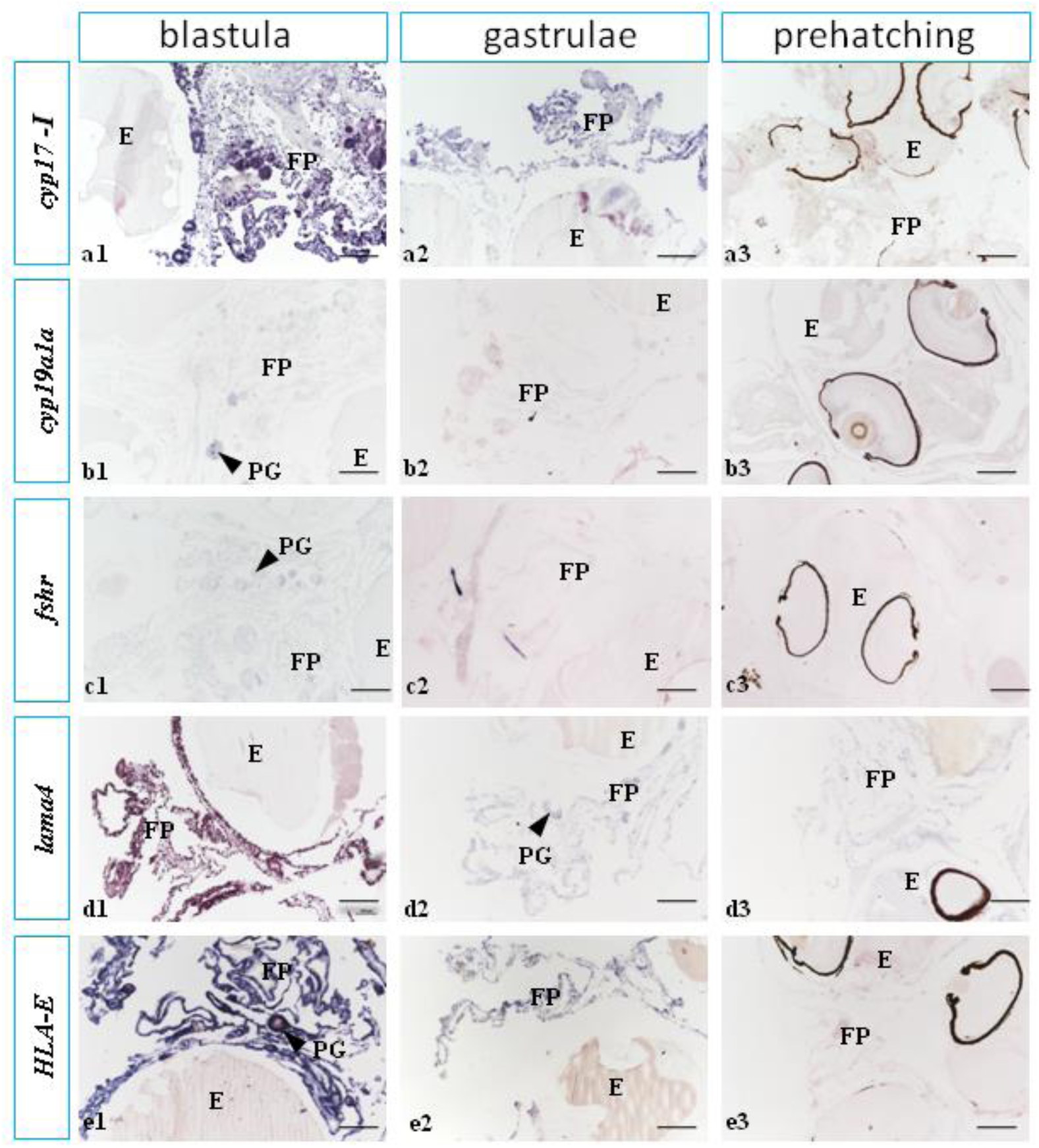
The expression of *cyp17-I, cyp19a1a, fshr, lama4* and *HLA-E* during pregnancy. *Cyp17-I* signals presented strong on the follicular placenta at blastula and gastrulae stage, and disappeared at prehatching stage (a1-a3). *Cyp19a1a* had no obvious signals during pregnancy (b1-b3). The expression of *fshr* during the gestation of black rockfish. *Fshr* had no obvious signals during gestation (c1-c3). *Lama4* expressed on the follicular placenta during the whole pregnancy period, the signals were strong at blastula and gastrulae stage, and became weak in the prehatching stage (d1-d3). *HLA-E* signals presented strong on the follicular placenta at blastula and gastrulae stage, and disappeared at prehatching stage (e1-e3). PG, follicles with primary growth; FP, follicular placenta; E, embryo. Scale bars, 200μm.

**Figure 6.**
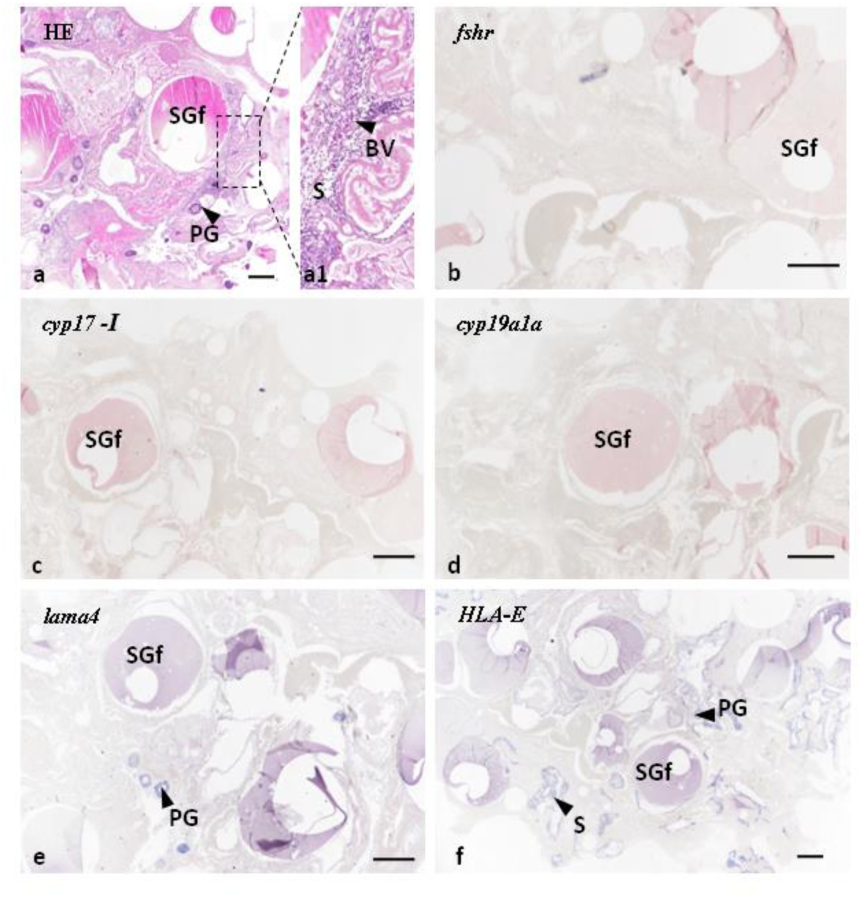
The morphogy and *cyp17-I, cyp19a1a, fshr, lama4* and *HLA-E* localization in the isolated female black rockfish ovary. No obvious follicular placenta structure was observed in the ovary, and only stromal cells, vascular structure and non-cellular structure around the unfertilized eggs (a). *Fshr, cyp17-I*, and *lama4* was not expressed on connection tissues around the oocytes (b-e), except *HLA-E* (f). PG, follicles with primary growth; SGf, full secondary growth; S, stromal cells; BV, blood vessel. Scale bars, 200μm.

## 3. Discussion

In black rockfish, we found a new type of folliculogenesis and placentation which is different from the other viviparous teleost. Vitellogenesis of black rockfish is similar to the turbot and other oviparous fish in general (*Lubzens et al., 2010*) (Fig. 7a, b). However, at SGe stage, the germinal epithelium gradually surrounded the SGe oocytes and formed a lot of individually developing follicles hanging on the ovigerous lamella with vascularized stalk-like structures attaching to the stroma, which guaranteed each follicle had opportunity contacting the spermatozoa as well as absorbing nutrition from ovary (Fig. 7b). After fertilization, we did not find the 2-cell structure outside the embryos by histology, suggesting follicular layers ruptured before fertilization, which was in line with the opinion of Bretschneider and Dewit (1947). After that, the stored spermatozoa fertilized the eggs, and no distinct boundary between the granulosa layer and theca layer were observed. The theca cells (with or without granulosa cells) proliferated rapidly and invaded into the surrounding connective tissue, becoming highly hypertrophied, extensively folded and highly vascularized, and quickly formed a microvillous placenta at blastula stage. The structure strongly resembles other teleost placenta structures and the portion of mammalian chorioallantoic placenta (*Grove and Wourms, 1994; Laksanawimol et al., 2006; Kwan et al., 2015*) (Fig. 7c).

**Figure 7.**
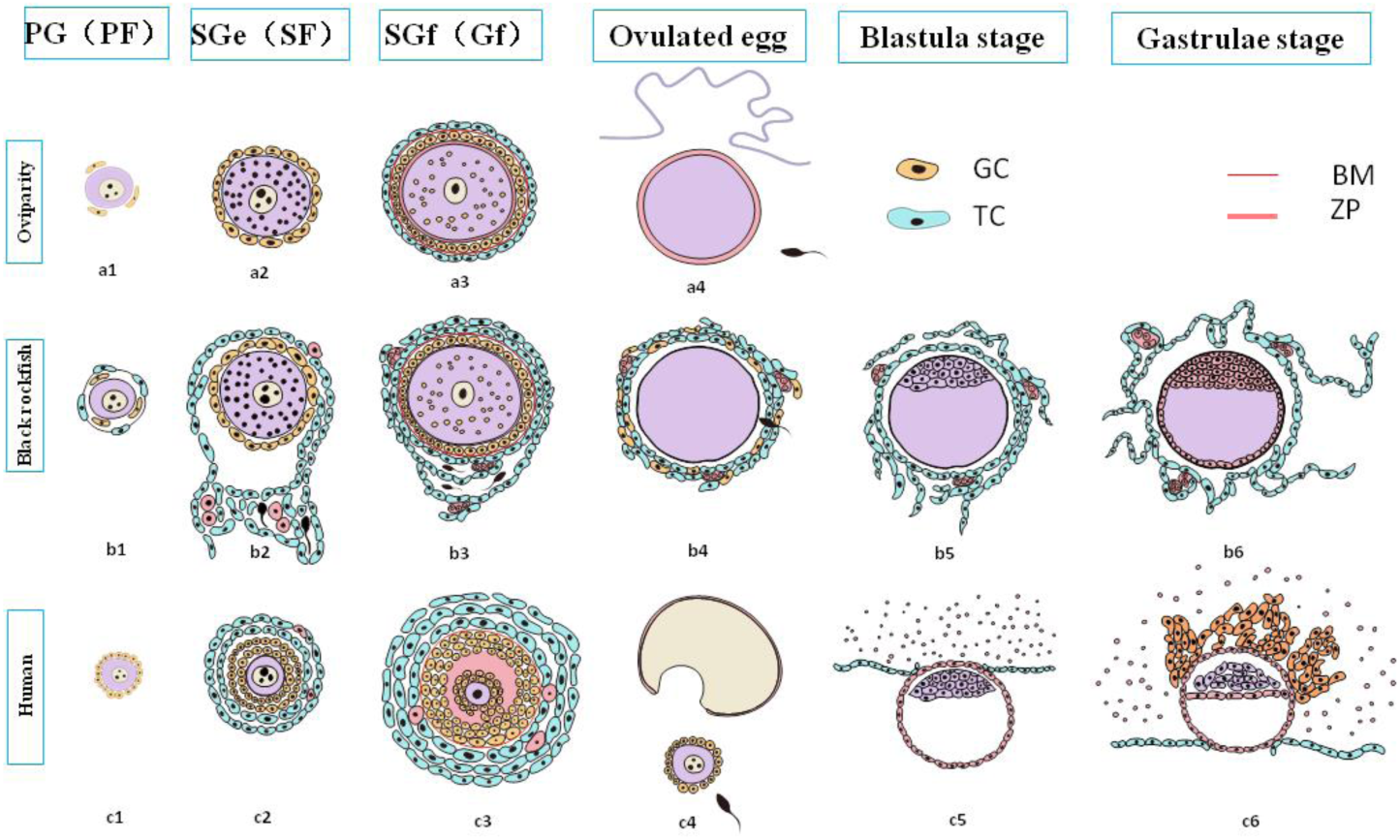
Cartoon illustrating the morphological difference during oogenesis and placentation among turbot (oviparity), black rockfish and human. Oogenesis and ovulation in turbot(oviparity) (a). After the follicles mature, the eggs are ovulated and fertilized in the water (a4). Oogenesis and placentation in black rockfish (b). After the follicles mature, the follicle layers rupture while the spermatozoa enter the micropyle (b4), then theca cells rapidly proliferate, migrate and invade outward forming the placenta (b5, b6). Oogenesis and placentation in human (c). After the follicles mature, the eggs are ovulated from the ovary to the fallopian tubes (c4), the sperm and egg unite to form a zygote (c5). Then the zygote travels down the fallopian tube and reaches the uterus. The morula becomes a blastocyst and implant into the uterine (c5, c6). PG, follicles with primary growth; SGe, early secondary growth; SGf, full secondary growth; PF, primary follicle; SF, secondary follicle; GF, graafian follicle; TC, theca cells; GC, granulosa cells; BM, basement membrane; ZP, zona pellucida.

Intriguingly, we also found some conserved genes derived from the mammals placentation expressed strongly during the early gestation in black rockfish. *Plgf*, a member of the vascular endothelial growth factor family (*Arroyo et al., 2008*), exclusively expressed in the early gestation in black rockfish. It can regulate vasculogenesis and angiogenesis of the placenta, and cause endothelial cell proliferation, migration, and tube formation (*Otrock et al., 2007; Tammela et al., 2005; Wallner et al., 2007*). *Plac8*, first recognized as a placenta-specific transcribed gene in mouse (*Galaviz-Hernandez et al., 2003*), also strongly expressed from stage V to early gestation period in black rockfish as well as in the follicular placenta of *P. retropinna* (*Guernsey et al., 2020*). *Plac8* has been found promoting trophoblast invasion and migration (*Mourtada-Maarabouni et al., 2013; Rogulski et al., 2005; Chang et al., 2018*). In addition, *tpbg* is prominently expressed in cleavage and blastula stage in black rockfish. *Tpbg* is abundantly expressed at the apical microvillus surface of the syncytiotrophoblast throughout gestation, but rarely expressed in other tissues (*Hole et al., 1988*). Besides that, TPBG is found to be released both from placental explants and perfused placenta, and sensitizes the maternal immune system (*Alam et al., 2017*). All of the above results indicated black rockfish shared the homology with genes of placentation in pregnant mammals, which further confirmed the follicular placenta structure existing during its gestation period.

Another interesting finding is that the ISH results from *cyp17-I* showed strong expression signals throughout the microvillous structure surrounding the embryos. Even before fertilization, the signals already exhibited obvious expansion especially in the stalk-like region. However, the *fshr* and *cyp19a1a* signals became weak at SGf stage and disappeared after fertilization, which also indicated the 2-cell structure breakdown, and granulosa cells lost the E2 synthesis function. Accordingly, the level of 11-KT progressively rose, peaked at blastula stage and remained a relative high level during the whole gestation period, while the E2 drastically decreased from stage IV and reached the bottom at blastula stage, then went back to a relatively high level. The high expression of *cyp19a1b* in gestation period might partially explained the high E2 level during the mid to late gestation period (*Kwon and Kim, 2013*). Similarly, progesterone also kept a high level from blastula stage, suggesting its important role in supporting pregnancy.

The changes of the three steroid hormone of black rockfish are similar to the previous studies (*Mu et al., 2013, Mori et al., 2003*), but we found the E2 rapidly declined to the bottom at blastula stage, while the 11-KT peaked at the same time. The asynchronous secretion of estrogen and androgen is different from the oviparous teleost, in which the E2 and 11-KT synchronously change with the oocyte development (*Kagawa, 2013*). Similar phenomenon has been reported in the prostate cancer (*Black et al., 2014*), in which CYP17A1 highly expressed and mediated intracellular androgens synthesis. Risk of aggressive prostate cancer was strongly inversely associated with estradiol: testosterone ratio (*Black et al., 2019*), and CYP17A1 is widely used as a target for the hormonal treatment of prostate cancer (*David et al., 2019*). The overexpression of *cyp17-I* at stage V led the unbalance between the 11-KT and E2, which might be the driven factor for the theca cells proliferation and invasion and form the microvillous placenta in black rockfish.

The site of gestation must be compartmentalized from the rest of the maternal tissues to maintain the appropriate environment for embryonic. HLA-G is important for the modulation of the maternal immune system during pregnancy, for it facilitates trophoblast invasion and fusion with maternal uterine arteries through inhibiting NK and T cell-mediated cell lysis (*Navarro et al., 1999; Rajagopalan and Long, 1999; Riteau et al., 2001a; Ishitani et al., 2003*). In this study, we identified *HLA-E* from black rockfish, which had the same conserved domains as the human. HLA-E and HLA-G all belong to the human MHC I genes (*Geraghty et al., 1987*, *1990*), and HLA-G was known as the specific molecular typical marker of extravillous trophoblasts (EVT) (*Ellis, 1990*; *Kovats et al., 1990*). HLA-E also plays a role in inhibiting natural killer cells by interacting with the CD94/NKG2A inhibitory receptor or activating CD94/NKG2C receptor (*Ashley et al., 2000; Braud et al., 1998*).

The results from both *ISH* and transcriptom showed that HLA-E was significantly expressed on the follicular placenta especially at SGf and blastula stage, which indicated *HLA-E* might play a similar role to *HLA-G*, faciliating the microvillous structure invasion during the early gestation stage. At the same time, we found the *lama4* also had a similar expression pattern to *HLA-E. Lama4* belongs to the laminin family, which is the basement membrane component that promotes cell adhesion and angiogenesis (*Givant-Horwitz et al., 2004*). LAMA4 was specifically localized in human first-trimester placental villi to syncytiotrophoblast cells and in the decidua to EVT cells, and promoted trophoblast invasion, migration and angiogenesis (*Shan et al., 2015*). We also detected the strong expression of *lama4* on the follicular placenta, suggesting the *Lama4* is one of crucial factors in cell invasion and angiogenesis in black rockfish. In black rockfish, both *HLA-E* and *lama4* were providing a microenvironment for placental cells proliferation, migration, invasion and signaling as reported in mammals (*Kim et al., 2014; Graubner et al., 2018*), which to some extent provides evidence for convergent evolution at molecular level on placentation in vertebrates.

In conclusion, we firstly demonstrated a new type of follicular placenta formed in Scorpaenidae, and unveiled the placentation was derived from the *cyp17-I* drastically strong expression leading to the unbalance between the 11-KT and E2. In addition, we found some highly conserved genes expressed in mammalian placenta were also significantly expressed in the black rockfish follicular placenta structure, suggesting the high convergence both in the fish and mammalian placenta evolution. This finding provided a new type of placentation pattern for viviparous teleost between the intrafollicular gestation and intraluminal gestation.

## 4. Materials and methods

### 4.1 Sample colletion

Females black rockfish were collected from September to May from Nanshan market, Qingdao, China. In addition, we isolated some females in the Shenghang Sci-tech Co, Ltd. (Weihai, Shangdong Province, China) before copulation, and collected gonad samples when other females developed to the middle of pregnancy. The turbot (*Scophthalmus maximus*) samples were obtained from March to July from Shenghang Sci-tech Co, Ltd. (Weihai, Shangdong Province, China).

Before collecting ovaries, all fish were anesthetized in tricaine methanesulfonate (MS-222, Sigma, St. Louis, MO). Half of each ovary was stored in liquid nitrogen for molecular experiments and transcriptome analysis, one quarter was fixed overnight in Bouin’s solution and preserved in 70% ethanol for histology observation. And the other quarter was fixed overnight in 4% paraformaldehyde (PFA) and preserved in 70% ethanol for ISH. Blood samples were collected from the caudal vein, settling in 4°C overnight and centrifuging at 16,000g for 10 minutes, and then stored at −80°C for hormones determination.

### 4.2 Total RNA extraction and cDNA synthesis

Total RNA was extracted from black rockfish ovaries using SPARK easy tissue/cell RNArapid extraction kit (SparkJade, China) following the manufacturer’s instructions. The RNA samples were determined by UV spectroscopy at 260 and 280 nm to measure concentration. The cDNA was synthesized by the PrimeScript™ RT reagent Kit with gDNA Eraser (Takara, Japan) and stored at −20 °C.

### 4.3 Histology

The histology followed the methods described by Yang et al. (2019) in our laboratory(. The fixed samples were dehydrated and embedded and then sliced, with a thickness of 5um (Leica 2235). After hematoxylin-eosin (H&E) staining, the morphological structures were observed under the microscope (NikonENi, Japan) at different stages.

### 4.4 *In situ* hybridization and the fluorescence *in situ* hybridization

The full length of the HLA-E sequence was obtained by our existing transcriptome data, the primers were shown in table 1. The synthesized cDNA was inserted into a pGEM-T Easy vector (Promega, Madison, WI) and the full length was verified by sequencing.

**Table 1.**
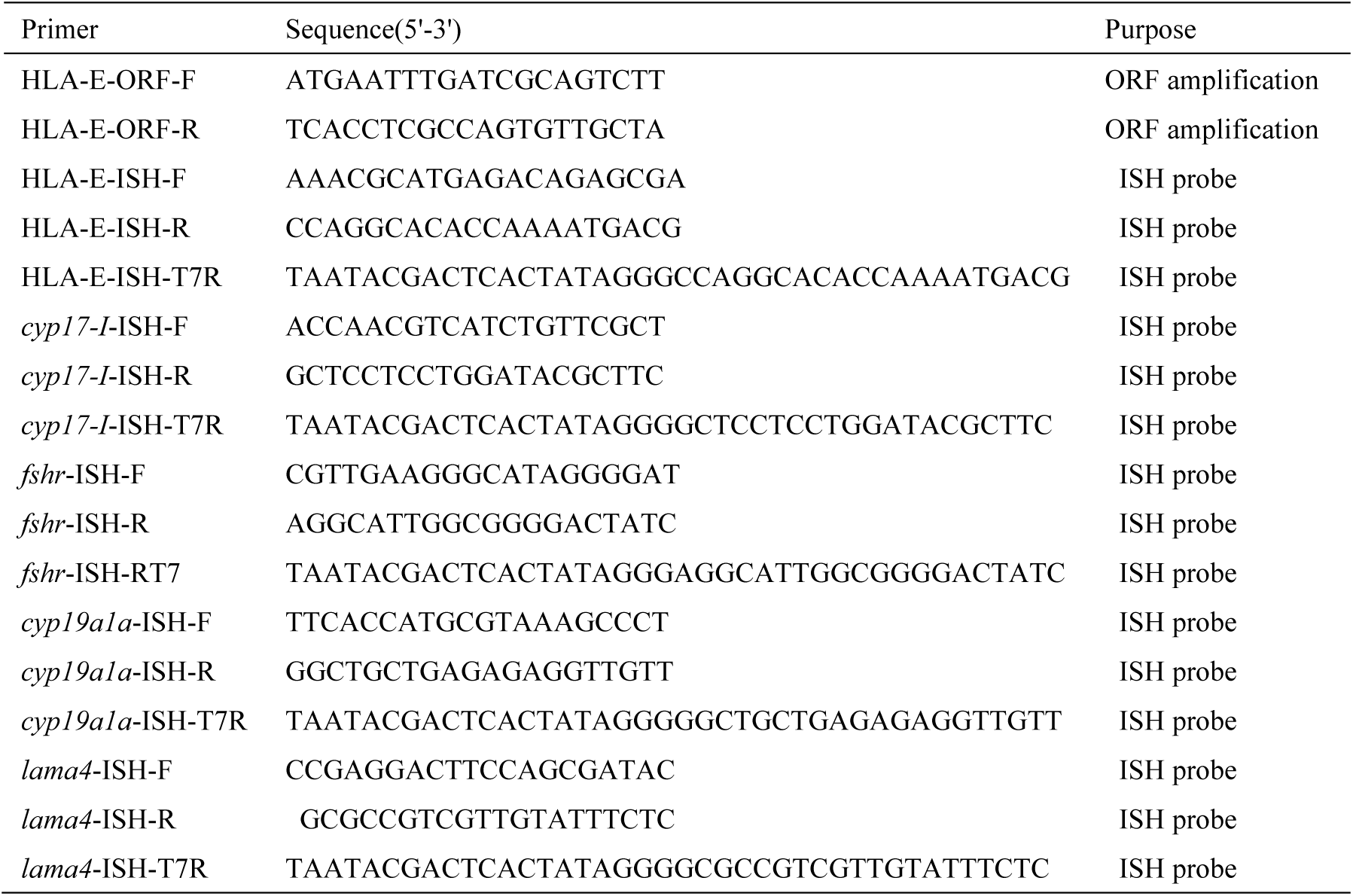
Primers and probes used for cloning and ISH

The full length of *cyp17-I, fshr* and *cyp19a1a* of black rockfish were carried out using the National Center for Biotechnology Information website(GenBank: ADV59774.2, AEJ33654, ACN39247), and the sequence of *lama4* was obtained by our existing transcriptome data.

The full length of *cyp17-I, fshr, cyp19a1a* and *lama4* of turbot were carried out using the National Center for Biotechnology Information website (GenBank: XM_035606144.1:142-1371, XM_035611916.1, XM_035627469.1, XM_035618004.1). Referring to a previous article (*Wang et al., 2017*),for each gene, primers F and R were used to amplify the cDNA fragment, and the product was used for the second round PCR using the primers F and R-T7 for generating antisense probe and primers R and F-T7 for generating sense probe used in ISH assays with the DIG RNA Labeling Kit (Roche, Mannheim, Germany),the primers were shown in table 1. After ISH, the samples were then stained with neutral eosin.

Two-color in the fluorescence *in situ* hybridization experiment was performed following the instructions of DIG RNA Labeling Kit (Roche, Mannheim, Germany). When synthesizing probe, *cyp17-I and cyp19a1a* were labelled with digoxin, and fluorescein, and detected by anti-dig and anti-fluorescein-POD antibodies, respectively.

### 4.5 Transcriptome analysis

Transcriptome analysis referred to a previous article (*Wang et al*,. *2018*). Twenty-four cDNA libraries (FII, FIII, FIV, FV, Cleavage, Blastula, Gastrulae, Prehatching) were constructed using total RNA from female ovaries at different development stage. The clean reads were assembled into non-redundant transcripts, which are then clustered into Unigenes. There were three biological repetitions for each stage.

### 4.6 Hormones determination

The E2, 11-KT, and P levels were tested by Iodine [125I] Radioimmunoassay (RIA) kits (Beijing North, China) respectively with the manufacturer’s instructions. The binding rate is highly specific with low cross-reactivity to other steroids, which was less than 0.1% to most circulating steroids.

### 4.7 Statistical analysis

The amino acid sequences of HLA-E of black rockfish was deduced using DNAMAN 8 software. The conserved domains of HLA-E genes in humans and black rockfish were predicted online through SMART(http://smart.embl-heidelberg.de/). Phylogenetic analysis was conducted with Mega7 software using the neighbor-joining method. The heat map was drawn with R software (3.5.3) based on the existing transcriptome data. FPKM (expected number of Fragments Per Kilobase of transcript sequence per Millions base pairs sequenced) was used to calculate the gene expression levels.

## Acknowledgement

This research was supported by National Key Research and Development Program (NO. 2018YFD0901205 2018, YFD0901204) and China Agriculture Research System (NO. CARS-47), and Major Science and Technology Innovation Projects 2019JZZY020710°

## Funding

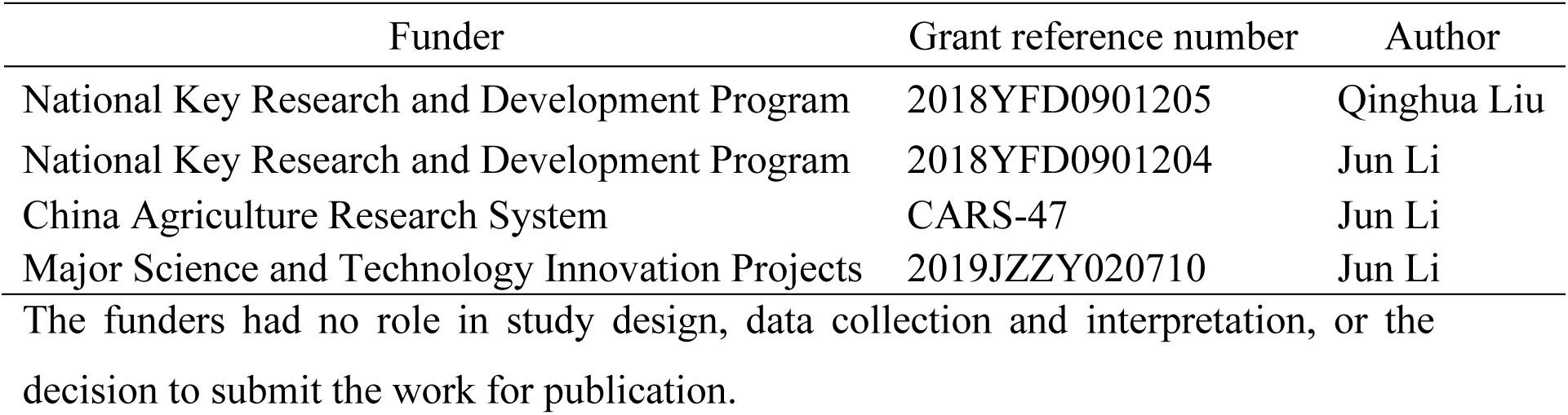

The funders had no role in study design, data collection and interpretation, or the decision to submit the work for publication.

## Author contributions

Xiaojie Xu, Conceptualization, Data curation, Formal analysis, Investigation, Methodology, Writing - original draft; Qinghua Liu, Conceptualization, Data curation, Formal analysis, Investigation, Methodology, Writing – review and editing, Funding acquisition, Supervision; Xueying Wang, Conceptualization, Formal analysis, Methodology; Xin Qi, Hormones level tests; Li Zhou, Methodology; Haoming Liu, fish culture; Jun Li, Supervision, Funding acquisition, Validation, Project administration

## Ethics

All experiments were performed in accordance with the relevant national and international guidelines and approved by the Institutional Animal Care and Use Committee, Institute of Oceanology, Chinese Academy of Sciences.

